# Distinct computational mechanisms of uncertainty processing explain opposing exploratory behaviors in anxiety and apathy

**DOI:** 10.1101/2024.06.04.597412

**Authors:** Xinyuan Yan, R. Becket Ebitz, Nicola Grissom, David P. Darrow, Alexander B. Herman

## Abstract

Decision-making in uncertain environments often leads to varied outcomes. Understanding how individuals interpret the causes of unexpected feedback is crucial for adaptive behavior and mental well-being. Uncertainty can be broadly categorized into two components: volatility and stochasticity. Volatility is about how quickly conditions change, impacting results. Stochasticity, on the other hand, refers to outcomes affected by random chance or “luck”. Understanding these factors enables individuals to have more effective environmental analysis and strategy implementation (explore or exploit) for future decisions. This study investigates how anxiety and apathy, two prevalent affective states, influence the perceptions of uncertainty and exploratory behavior. Participants (N = 1001) completed a restless three-armed bandit task that was analyzed using latent state models. Anxious individuals perceived uncertainty as more volatile, leading to *increased* exploration and learning rates, especially after reward omission. Conversely, apathetic individuals viewed uncertainty as more stochastic, resulting in *decreased* exploration and learning rates. The perceived volatility-to-stochasticity ratio mediated the anxiety-exploration relationship post-adverse outcomes. Dimensionality reduction showed exploration and uncertainty estimation to be distinct but related latent factors shaping a manifold of adaptive behavior that is modulated by anxiety and apathy. These findings reveal distinct computational mechanisms for how anxiety and apathy influence decision-making, providing a framework for understanding cognitive and affective processes in neuropsychiatric disorders.

## Introduction

Life is filled with unexpected challenges. How individuals interpret the causes of undesirable outcomes, such as investment failures, career plateaus, or bad weather, in uncertain environments shapes their subsequent actions (1). When people attribute changes in outcomes to environmental volatility (the speed at which the environment is changing), they may be motivated to explore more, seeking additional information and altering their behavior. In contrast, attributing adverse outcomes to mere chance or “bad luck” (stochasticity) may decrease the motivation to explore, leading some individuals to persist with their existing strategies (2).

The response to environmental uncertainty likely interacts with individuals’ affective states in a bidirectional manner. Attributing adverse outcomes to stochasticity may lead individuals to stick to previous behaviors, potentially protecting them from hurtful feedback through additional interaction with the world. However, this approach may also dampen an individual’s ability to adapt to a changing environment, potentially reinforcing a negative cycle and leading to apathy and depression. Conversely, perceiving sources of negative outcome as volatile may motivate individuals to learn more about the world and reduce uncertainty, though this may also increase the chances of experiencing more adverse outcomes and potentially worsening negative feelings such as anxiety.

Reciprocally, how individuals perceive and respond to environmental uncertainty can be influenced by underlying affective states (3). Apathy, characterized by a lack of motivation and goal-directed behavior (4, 5), is an affective state associated with imprecise beliefs about action outcomes (6) and a tendency to persist with previous choices rather than explore (7). This suggests that apathetic individuals may view outcomes as primarily stochastic, attributing events more to chance than controllable variables. This bias could discourage exploration and potentially reinforce a cycle of failure and helplessness (8).

In contrast, anxiety, marked by excessive worry and a heightened perception of potential threats (9, 10) and uncertainty (11), may lead individuals to overestimate environmental volatility. Consequently, anxious individuals could be driven to seek new information to update their beliefs and reduce uncertainty (12). However, research on the link between anxiety and exploration has yielded mixed findings, with some studies showing increased exploration to mitigate uncertainty (13, 14) and others showing reduced exploration to avoid unpredictable feedback under high anxiety (15, 16). Notably, apathy and anxiety often coexist in clinical populations, such as Alzheimer’s (17), Parkinson’s disease (18), and depression (19), despite having distinct neural representations (20, 21).

Building on these findings, we propose three fundamental questions to further elucidate the relationship between affective states and decision-making under uncertainty. First, we aim to investigate whether apathy and anxiety exhibit distinct behavioral patterns when individuals are faced with uncertain situations. Second, we seek to examine how individual differences in levels of apathy and anxiety are associated with perceptions of different types of uncertainty, specifically volatility and stochasticity. Finally, we intend to explore how perceived volatility influences exploratory behavior during decision-making processes.

We posit two competing hypotheses:

1. Apathetic individuals manifest less exploration, while anxious individuals engage in more exploration. Apathetic individuals weigh stochasticity over volatility and explore less, while anxious individuals overestimate volatility but explore more to reduce their uncertainty. This result would be consistent with previous findings suggesting that the two affective states have distinct neural substrates (20, 22).
2. Both apathetic and anxious individuals engage in less exploratory behavior but through different computational mechanisms. Apathetic individuals weigh stochasticity over volatility and explore less, while anxious individuals overestimate volatility, leading to a sense that their actions cannot track or learn from the environment, ultimately leading to exploitation. This may provide a computational account for learned helplessness (23) and the co-occurrence of apathy and anxiety in various clinical populations, such as Parkinson’s and Alzheimer’s diseases.

To address these questions, we employed a restless three-armed bandit task (Figure 1A), a well-established paradigm for capturing adaptive learning in volatile environments(24). We adopted Hidden Markov Model (HMM) to obtain the likelihood of individuals switching between exploitation and exploration states (25, 26). To further investigate how volatility and stochasticity modulate exploration, we utilized a Kalman filter model, which can dissociate two distinct sources of noise, volatility (process noise variance) and stochasticity (observation noise variance), during inference (27). Together, these methods offer a comprehensive view of the cognitive mechanisms underlying exploratory behavior and the manifestation of anxiety and apathy.

**Figure 1.**
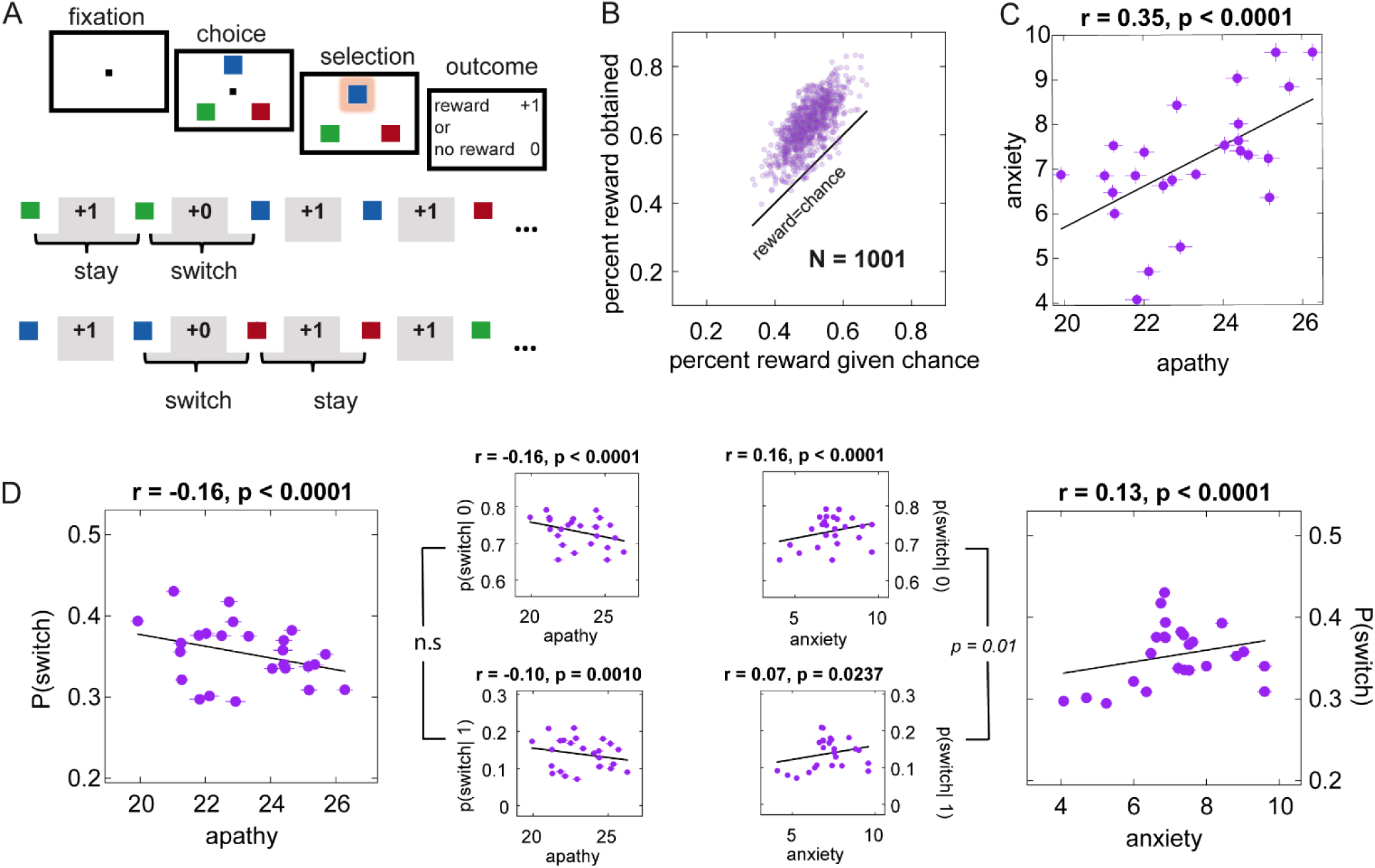
Three-armed restless bandit task and distinct behavioral patterns associated with apathy and anxiety. (A) Three-armed restless bandit task. Participants chose one option from among the three targets to receive reward or non-reward feedback. Each target was associated with a hidden reward probability that randomly and independently changed throughout the task. The lower panel indicates the example choice and reward sequence and the definition of stay and switch. Specifically, stay was defined as choosing the same target as in the previous trial, while switch was defined as choosing a different target. “+1” denotes reward feedback, and “+0” denotes reward omission. (B) Most participants earned more rewards than expected by chance (C) Apathy and anxiety correlated positively. (D) Apathy correlated negatively with switch behaviors, while anxiety correlated positively with switch behaviors. Anxious individuals were more sensitive to undesired feedback (no reward) and exhibited more switch behaviors compared to reward feedback. (Panels in Figure 1C and 1D utilize binned correlation plots [25 quantile bins based on the x-axis], with lines representing the standard error (S.E.). N.B. that these may be smaller than the symbol. Statistical analyses were performed on raw data.)

## Results

We recruited a large gender-balanced online sample consisting of 1,001 adults. The participants, ranging in age from 18 to 54 years (mean ± SD = 28.446 ± 10.354 years; gender: 493 female), performed a restless three-armed bandit task, as depicted in Figure 1A. During this task, participants selected among three playing card images, with each card representing a different option. They made their selections by moving their cursor over their chosen card. The probability of receiving a reward from each card deck varied randomly over time. After each choice, feedback was displayed on the screen indicating whether a reward was received. Participants also completed symptom surveys assessing levels of anxiety and apathy (details in Methods and SI Section 1, Table S1). We defined the trial as a *switch* trial if the chosen option was different from the last trial, and a *stay* trial if the choice was the same as the last trial.

### Apathy and anxiety predicted distinct exploratory behaviors

We first evaluated the performance by comparing the total number of rewarded trials each participant experienced against the number expected by chance. Out of the 1001 participants, 985 accrued more rewarded trials than would be statistically expected by chance, suggesting significant effectiveness in their decision-making strategies (Figure 1B). As expected, anxiety and apathy showed a significant positive correlation (r = 0.35, *p*<10^-29^, Figure 1C), which is consistent with previous findings on their co-occurrence (17).

To investigate the relationship between apathy and the percentage of switch behaviors (P(switch)), as well as anxiety and P(switch), we conducted partial correlations between apathy and exploration while controlling for anxiety, and between anxiety and exploration while controlling for apathy. We found that apathy negatively predicted P(switch) (r = −0.16, *p*<0.001) regardless of feedback type (reward or no-reward), while anxiety positively correlated with P(switch) (r = 0.13, *p*<0.001). Intriguingly, the relationship between anxiety and switch behaviors was greater after non-reward feedback (r = 0.16, *p*<0.001) compared to reward feedback (r = 0.07, *p*=0.024) (their difference, z-score = 2.40, *p*=0.01). Though co-existing in this population, these two affective states predicted distinct switch behaviors under uncertainty (Figure 1D). The stronger relationship between anxiety and P(switch) after undesirable feedback indicates that highly anxious individuals are more sensitive to negative feedback, which may lead them to disengage.

Next, we fitted the behavior with a Hidden Markov Model (HMM) to decode the hidden states, “*explore*”, and “*exploit*” (Figure 2A) (24, 25, 28, 29). Each arm is associated with a hidden reward probability that randomly and independently changes throughout the task (Figure 2A). In our study, exploration and exploitation states are considered hidden states underlying the observed choices, such as switching between decks or repeatedly choosing from the same deck. We calculated the percentage of explore states, i.e., P(explore). Consistently, apathy correlated negatively with P(explore) (r = −0.17, *p*<0.001), while anxiety positively correlated with (P(explore)) (r = 0.11, *p*=0.003) as well as the percentage of exploration after reward omission(P(explore|0) (r = 0.13, *p*<0.001) (Figure 2C).

**Figure 2.**
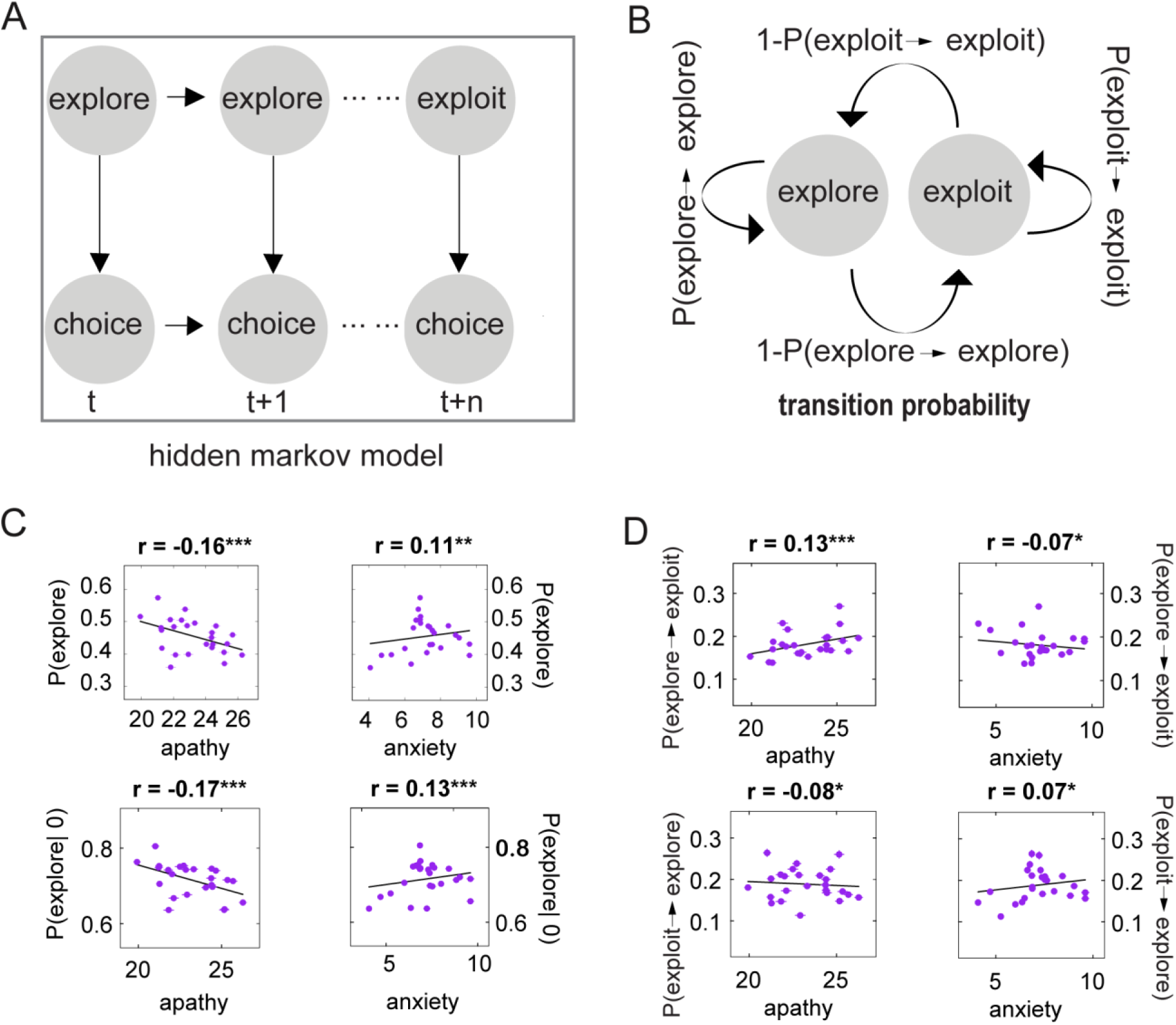
Apathy and anxiety have opposing relationships with exploration and explore and exploit state dynamics. (A) Unrolled structure of the hidden Markov model (HMM) used to infer the explore and exploit states’ underlying behavior. (B) The transition probabilities within and between states in the HMM. (C) The probability of exploration, plotted as a function of apathy (top left) and anxiety (top right). The probability of exploration following a reward omission is plotted as a function of apathy (bottom left) and anxiety (bottom right). (D) The transition probability from explore to exploit, plotted as a function of apathy (top left) and anxiety (top right); the transition probability from exploit to explore plotted as a function of apathy (bottom left) and anxiety (bottom right). (Panels in Figure 2C and 2D utilize binned correlation plots [25 quantile bins based on the x-axis], with lines representing the standard error (S.E.). N.B. that these may be smaller than the symbol. Statistical analyses were performed on raw data). * *p* < 0.05, ** *p* <0.01, *** *p* < 0.001. All p-values remained significant after FDR *p*<0.05 correction.

In addition to the overall frequency with which hidden states occur, examining the transitions between these states can further illuminate the dynamics of decision-making. Therefore, we investigated how apathy and anxiety manifest in the transition probability (Figure 2B) between explore and exploit. As predicted, apathy had a positive correlation with the transition probability from *explore* to *exploit* (r = 0.13, *p*<0.001) but a negative correlation with the transition probability from *exploit* to *explore* (r = −0.08, *p*=0.011). In contrast, anxiety had a negative correlation with the transition probability from *explore* to *exploit* (r = −0.07, *p*=0.035) but a positive correlation with the transition probability from *exploit* to *explore* (r = 0.07, *p*<0.022) (Figure 2D). All significant results reported in the study survived False Discovery Rate (FDR, *p*<0.05) correction.

### Apathy and anxiety are associated with distinct computational processes underlying exploration

We then asked whether differing perceptions of the environment might explain the distinct patterns of exploration predicted by apathy and anxiety we observed.

To address this question, we utilized a Kalman filter model (Figure 3A), which can dissociate sources of uncertainty into perceived volatility and stochasticity (27). Kalman filter (KF) models have been widely applied in psychology and neuroscience to study various aspects of learning and decision-making (30, 31) (for more detailed information about the model, please refer to the Method section).

**Figure 3.**
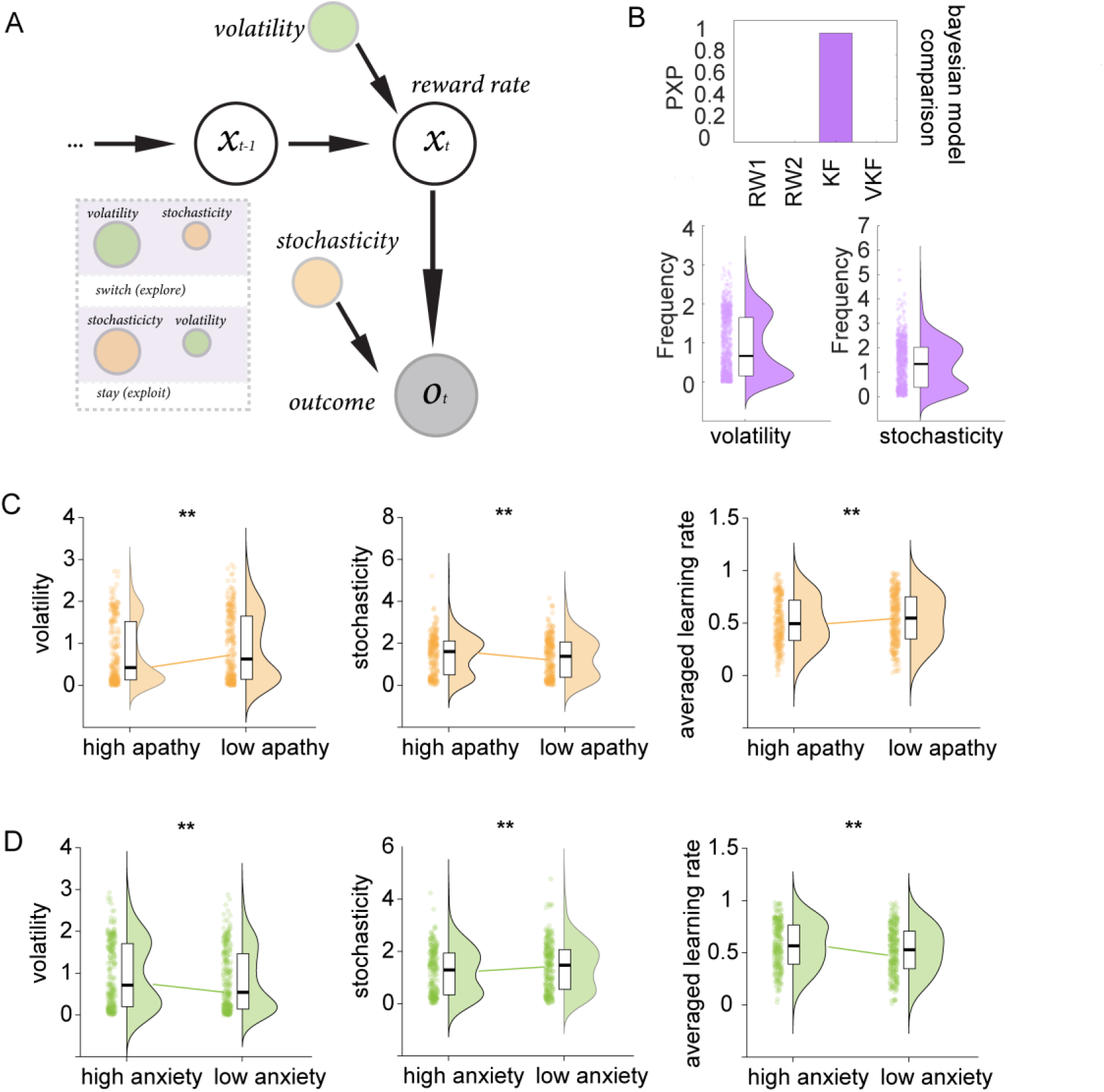
Apathy and anxiety have opposing relationships with volatility and stochasticity. (A) The schematic of the Kalman filter model used in our analysis. The diagram illustrates how this model can differentiate between volatility (process noise variance) and stochasticity (observation noise variance), providing insights into the underlying decision-making processes. (B) Bayesian model comparison and the distribution of volatility, stochasticity (C) Highly apathetic individuals overestimated stochasticity but underestimated the volatility, resulting in a lower learning rate. (D) In contrast, highly anxious individuals overestimated volatility but underestimated stochasticity, resulting in a higher learning rate. \**p* < 0.05, ** *p* <0.01, *** *p* < 0.001. All p-values remained significant after FDR *p*<0.05 correction.

We also fitted the behavioral data to alternative models including volatile Kalman filter (VKF) (27), Rescorla-Wagner models single (RW1) (32) and dual learning rates (RW2) to weigh positive and negative learning rates (33). We employed Hierarchical Bayesian inference (HBI) to fit models to choice data (34). Further, we used Bayesian model selection (BMS) and protected exceedance probability (PXP) to select the winning model (Figure 3B). The Kalman filter served as the best model for our population, and we examined the resulting distribution of volatility and stochasticity (Figure 3B).

We first conducted correlation analyses using all data points. Specifically, we found that apathy was positively correlated with stochasticity (r = 0.08, *p*=0.013) but negatively correlated with volatility (r = −0.08, *p*=0.008). Conversely, anxiety showed a negative correlation with stochasticity (r = −0.12, *p*=0.001) and a positive correlation with volatility (r = 0.12, *p*=0.002). These correlations highlight the distinct cognitive biases associated with apathy and anxiety in processing environmental uncertainties.

To clearly illustrate and confirm the findings, we categorized participants into distinct groups based on their apathy and anxiety levels. For apathy, we identified the high apathy group (N = 223) as those scoring in the top 25% on the Apathy Motivation Index (AMI), which assesses apathy in behavioral and social domains (35). Conversely, the low apathy group (N = 251) comprised individuals scoring in the bottom 25% of apathy scores. Similarly, for anxiety, the high anxiety group (N = 228) included participants within the top 25% of scores on the GAD-7 scale (36), while the low anxiety group (N = 250) consisted of those in the bottom 25%. These classifications allowed for a direct comparison of behaviors and traits between individuals with varying degrees of apathy and anxiety.

We conducted linear regression analyses using volatility and stochasticity as the dependent variables with the high versus low anxiety and apathy groups as predictors. The Methods section provides details of the regression model specifications.

As hypothesized, apathetic individuals overestimated stochasticity (t(471) = 3.06, *p*=0.002) and underestimated the volatility compared to those with low apathy (t(471) = −3.24, *p*=0.001)(Figure 3C). Consequently, apathetic individuals exhibited a lower learning rate than their low apathy counterparts (t(471) = −3.11, *p*=0.002).

In contrast, individuals with high anxiety levels tended to overestimate volatility (t(475) = 2.84, *p*=0.004) and underestimate stochasticity compared to those with low anxiety (t(475) = −3.04, *p*=0.002), resulting in a higher learning rate among the high anxiety group (t(475) = 3.21, *p*=0.001) (Figure 3D). Furthermore, comparisons showed that anxious individuals had higher volatility estimates than those with high apathy (t(449) = 2.75, *p*=0.006), whereas apathetic individuals had higher stochasticity estimates than their anxiety counterparts (t(449) = −3.01, *p*=0.002) (SI Section 2, Figure S1).

### The ratio of volatility to stochasticity distinguished apathy and anxiety

To clarify the differential impacts of apathy and anxiety on decision-making under uncertainty, we computed the ratio of volatility to stochasticity, *ν*/*s* to represent the balance between these two types of uncertainties. A higher *ν*/*s* indicates a perception of greater volatility relative to stochasticity, while a lower ratio suggests a perception of more stochasticity relative to volatility. We applied a logarithmic transformation to the ratio to manage extreme values (e.g. cases where individuals might perceive very high volatility but very low stochasticity).

Consistently, our findings reveal a clear distinction: *ν*/*s* correlated negatively with apathy (r = - 0.08, *p*=0.010) but positively with anxiety (r = 0.13, *p*<0.001) (Figure 4A).

**Figure 4.**
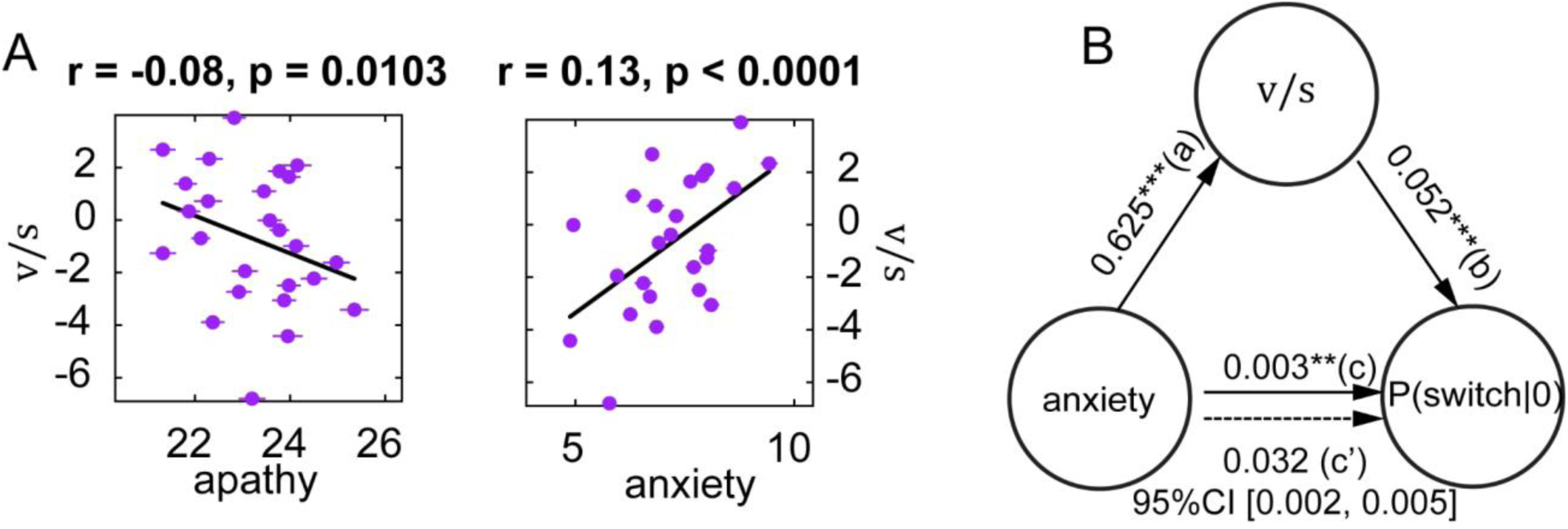
Distinctions in apathy and anxiety on the ratio of volatility to stochasticity and its mediation effect. (A) The ratio of volatility to stochasticity, plotted as a function of apathy (left) and anxiety positively (right). (B) Mediation analysis, showing the mediating effect of the ratio of volatility to stochasticity on the relationship between anxiety and switch behavior after reward omission. (Panels in Figure 4A utilize binned correlation plots [25 quantile bins based on the x-axis], with lines representing the standard error (S.E.). N.B. that these may be smaller than the symbol. Statistical analyses were performed on raw data). * *p* < 0.05, ** *p* <0.01, *** *p* < 0.001. All p-values remained significant after FDR *p*<0.05 correction).

### The ratio of volatility and stochasticity mediated the relationship between anxiety and the exploration after negative feedback

To determine whether individual differences in the perception of uncertainty explain the relationship between exploratory behavior and affect, we conducted a mediation analysis with anxiety, switching after reward omission (P(switch | 0)), and *ν*/*s*. The results demonstrate that the relationship between anxiety and the tendency to switch after receiving no reward is significantly mediated by *ν*/*s* (Figure 4B). This mediation was also significant for the analogous HMM model-based measures (see SI Section 3, Figure S2). No significant mediation effect was found for apathy, however, reinforcing the unique pathways through which anxiety influences exploratory behavior. These results explain why individuals with higher anxiety might explore more after negative feedback, driven by an overweighting of perceived volatility relative to stochasticity as a strategy to reduce uncertainty and manage risks.

### A low dimensional manifold unifies exploration, perceptions of uncertainty and affective state

The HMM state-model of exploration-exploitation and the Kalman filter process model of uncertainty estimation represent complementary ways of understanding adaptive behavior that our mediation results suggest are intrinsically related. We hypothesized that a latent structure underlying adaptive behavior on this task might unify these descriptions of behavior. We utilized advanced dimensionality reduction methods to uncover such a latent structure in the raw task behavior.

First, we formatted each participant’s trial-by-trial task data into sequences of choices to stay (repeat the choice on the last trial) or switch (choose a different option) and reward outcome for two consecutive trials ({choice_t-1_, outcome_t-1_, choice_t_}, Figure 5A and 5B). The behavioral data for each participant was then transformed into counts for each of these eight unique sequences. Then we applied Uniform Manifold Approximation and Projection (UMAP) (37), a computationally efficient algorithm that can preserve both the local and global distances between data points in high-dimensional space, to learn the two-dimensional manifold underlying the eight-dimensional behavioral data (Figure 5C, see Methods for more algorithm details).

**Figure 5.**
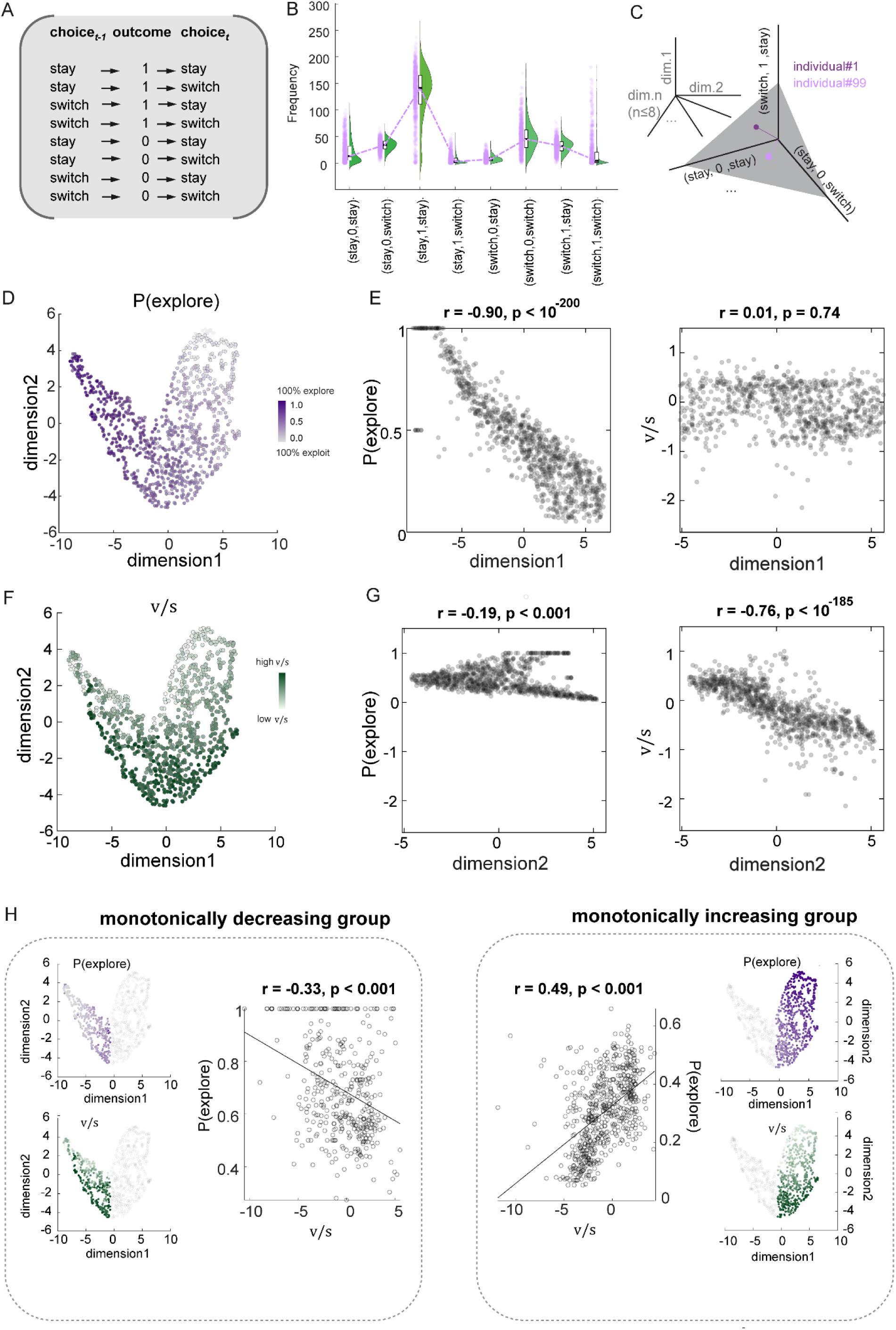
Visualizing the complex relationships in decision-making through low-dimensional space. (A) All possible sequences of choices and rewards that participants could make during the experiment (B) The frequency distribution of individual decision-making patterns. The black line in the box plot represents each pattern’s mean value, highlighting participants’ typical behaviors (C) Schematic high-dimensional space of participants’ decision-making pattern (D) The two-dimensional space representation of exploration by using the Uniform Manifold Approximation and Projection (UMAP) (Different dimensionality reduction methods such as principal component analysis (PCA), and t-distributed Stochastic Neighbor Embedding (t-SNE) lead to a similar space) (E) Dimension 1 exclusively represents P(explore) but does not represent the ratio of volatility to stochasticity (F) The two-dimensional space representation of the ratio of volatility to stochasticity behavior by using UMAP (Different dimensionality reduction methods such as principal component analysis (PCA), and t-distributed Stochastic Neighbor Embedding (t-SNE) lead to a similar space) (G) Dimension 2 mainly represents the ratio of volatility to stochasticity but not P(explore) (H) The manifold has been separately dissociated into the monotonically decreasing group (the most left panel) and monotonically increasing group (the most right panel). The monotonically decreasing group was associated with a relatively higher anxiety level than the monotonically increasing group, while the apathy level was significantly lower than the monotonically increasing group. Within the monotonically decreasing group (left part), a higher volatility to stochasticity ratio leads to decreased exploration. In contrast, within the monotonically increasing group (right part), a higher volatility to stochasticity ratio encouraged higher exploration. This exploration serves as a coping strategy to relieve anxious feelings in the environment. All p-values remained significant after FDR *p*<0.05 correction.

Including additional reward history and applying other dimensionality reduction methods like principle component analysis (PCA), and t-distributed Stochastic Neighbor Embedding (t-SNE) did not change the results (SI Section 4, Figure S3, Table S2).

Our analysis using UMAP revealed distinct correlations within the derived dimensions. Specifically, the dimension 1 score (the horizontal axis) exhibited a very strong significant negative correlation with exploratory behavior (P(explore) (r = −0.90, *p*<10^-200^), but it showed no significant relationship with the ratio of volatility to stochasticity (*ν*/*s*)(r = 0.01, *p*=0.74) (Figure 5D, 5E). In contrast, the dimension 2 score (the vertical axis) demonstrated a strong negative correlation with *ν*/*s* (r = −0.76, *p*<10^-185^), which was significantly more pronounced than its correlation with P(explore) (r = −0.19, *p*<10^-10^) (Figure 5F, 5G). This suggests that dimension 1 primarily represents exploratory behavior, while dimension 2 primarily reflects the computational factors: volatility and stochasticity (i.e., volatility and stochasticity).

Further, both dimensions also showed correlations with affective states: the dimension 1 score was positively correlated with apathy (r= 0.14, *p*<0.001), and negatively correlated with anxiety (r = - 0.11, *p*<0.001). Similarly, the dimension 2 score had a positive correlation with apathy (r = 0.097, *p*=0.002) and a negative correlation with anxiety (r = −0.088, *p*=0.004).

It is worth noting that we only found linear relationships between apathy, anxiety, and exploration, as well as between these affective states and the ratio of volatility to stochasticity (our analysis using higher order effects among these variables did not yield significant results, more details can be found in SI Section 5, Table S3).

To delve deeper into how these factors interact in the low-dimensional space defined by UMAP, we divided the data manifold into two groups based on the dimension1 score: a monotonically decreasing group (left part, dimension 1 score < −0.671, *N*=390) and a monotonically increasing group (right part, dimension 1 score> −0.671, *N*=611). The methodology used to identify the turning point (dimension 1 score = −0.671) that differentiates the monotonically decreasing group from the monotonically increasing group is detailed in the Methods section. The analysis revealed that the monotonically decreasing group had relatively higher levels of anxiety compared to the monotonically increasing group (t(999)=2.08, *p* =0.037), while their apathy levels were significantly lower (t(999)= −3.56, *p*=0.0003). This segmentation allows us to further explore and understand the complex interplay between affective states, computational parameters, and exploratory behaviors within a structured, low-dimensional framework. Notably, individuals in the left monotonically decreasing group, characterized by high anxiety and low apathy, generally perceive higher volatility relative to stochasticity and demonstrate greater exploratory behavior compared to those in the right monotonically increasing group. Within the decreasing group, higher perceived volatility correlates with reduced exploration. Conversely, in the increasing group, an increased perception of volatility tends to stimulate more exploratory actions. These results suggest that while severe anxiety might suppress exploration due to overwhelming uncertainty, moderate anxiety in the general population can promote exploration as a coping mechanism to gather information and reduce anxiety symptoms.

We now address our final research question; what is the relationship between volatility and exploratory behavior? Considering the parabolic relationship between the manifold dimension reflecting exploration and the dimension representing *ν*/*s*, we hypothesized that the relationship between exploration and *ν*/*s* might be quadratic.

To test this hypothesis, we constructed a regression model as follows:

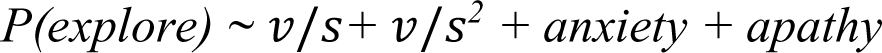

The results revealed that both the linear and quadratic terms are significant (linear term, coefficient = 0.02, SE = 0.003, t(996)=5.83, *p*<10^-9^; quadratic term, coefficient = 0.005, SE = 7.86×10^-4^, t(996)=6.59, *p*<10^-9^), indicating a complex, non-linear relationship between the ratio of volatility to stochasticity and exploration (see SI Section 6, Figure S4), which was consistent with the manifold representation.

## Discussion

We found that apathy and anxiety predicted opposing patterns of exploratory behavior, which were explained partly by differing perceptions of uncertainty. Anxiety was associated with increased exploration after reward omission and greater volatility estimation: the attribution of uncertainty to a rapidly changing (but still learnable) environment. Apathy, in contrast, predicted decreased exploration and higher stochasticity estimation: the perception of uncontrollable randomness. Following a dimensionality reduction of the raw behavioral data, exploration and perceptions of uncertainty emerged as the dimensions of an underlying latent structure that unified the different model approaches and the affective states. These findings elucidate the complex interplay between cognitive assessments of uncertainty, affective states, and decision-making processes, offering several key insights into adaptive and maladaptive behaviors under uncertainty.

The distinct patterns of exploratory behavior observed in anxious and apathetic individuals highlight the role of affective states in shaping responses to uncertainty. Anxious individuals, who generally display a heightened sensitivity to potential threats and environmental changes, exhibited a bias toward perceiving greater volatility and exploring more after negative outcomes. Our mediation analysis revealed that the perception of volatility relative to stochasticity partially mediates the relationship between anxiety and exploratory behavior after reward omission. This finding is consistent with previous results (38) and offers a mechanistic explanation for why anxious individuals in a healthy population might choose to explore more after receiving negative feedback. The perceived overweighting of volatility relative to stochasticity may drive these individuals to seek more information, potentially as a strategy to reduce uncertainty and manage perceived risks more effectively (39). Although such a strategy may be beneficial for adaptation in genuinely volatile environments, it may also contribute to excessive worry and stress, especially if the perceived level of volatility exceeds actual environmental volatility (9, 11). Consequently, anxious individuals may find themselves in a prolonged state of heightened arousal and uncertainty, leading to suboptimal decision-making and diminished well-being.

On the other hand, apathetic individuals, who generally exhibit diminished motivation and responsiveness (40), tended to attribute outcomes more to stochasticity in our study. This perception might underlie their reduced exploratory behavior, reflecting a disengagement from active learning and adaptation. If outcomes seem random and beyond our control, expending energy to explore may seem futile, and focusing on what we know seems rational. While this approach may conserve energy, the inflexibility can perpetuate a cycle of disengagement and maintain apathetic symptoms (41, 42). Apathetic individuals may fail to recognize the potential benefits of exploration and remain stuck in suboptimal decision-making patterns, further reinforcing their disengagement from the environment (4).

The dimensionality reduction of the behavioral sequence data using UMAP allowed us to examine the relationship between exploration and the estimation of volatility and stochasticity. Despite the intuitive connection between these two behavior models, their relationship has not been directly examined. Our results showed that exploration and uncertainty estimation related closely to the two axes of a parabolic latent structure of adaptive behavior. As a result, both model-based metrics were necessary to characterize the spectrum of individual differences fully.

Segmenting the data on the manifold further illuminated the fine-grained interplay between affective states and exploratory behavior. Individuals with relatively higher anxiety and lower apathy (the monotonically decreasing group) generally weighted volatility more and demonstrated greater exploratory behavior compared to those with lower anxiety and higher apathy (the monotonically increasing group). However, *within* these groups formed on the manifold, individuals exhibited opposing relationships between uncertainty and exploration. In the higher anxiety group, perceived volatility correlates inversely with exploration. However, in the lower anxiety group, increased volatility perception predicts greater exploration.

These results reconcile previously inconsistent findings regarding exploratory behavior in individuals with anxiety, with some studies showing more exploitative behavior (15, 16), and others finding that higher anxiety predicts more exploratory behaviors (13, 14). The relationship between perceived volatility and exploration is modulated by the degree of anxiety, with more severe anxiety potentially suppressing exploration as a form of avoidance. Conversely, moderate anxiety may drive exploration to gather information and reduce uncertainty, potentially easing discomfort. This dual response to perceived volatility underscores the complex interplay between anxiety levels, environmental perceptions, and behavioral strategies in managing emotional responses.

Our findings have implications for personalized behavioral interventions in mental health. For anxious individuals, therapies focusing on recalibrating volatility perceptions and improving uncertainty management may reduce worry and enhance decision-making (43, 44). Encouraging longer-term information integration could also benefit anxiety management (38). For apathetic individuals, strengthening perceived control and action efficacy may counteract stochasticity attribution. Incorporating these strategies into existing therapies like Behavioral Activation and Motivational Interviewing (45) could promote balanced environmental perceptions and exploration.

Another important clinical implication involves using an individual’s position on the behavioral manifold (Figure 5) to predict how their behavior might change in response to treatment based on their symptoms. For example, a patient positioned in the upper left quadrant before treatment may exhibit higher anxiety, lower apathy, and increased exploratory behavior. During and after treatment, monitoring these behavioral shifts may allow us to infer changes in their affective states or symptoms based on their new manifold position. To develop such a tool, several questions remain: Do changes within an individual follow a predictable trajectory on this manifold? Do clinical populations conform to the same manifold, or do they deviate, projecting into the larger, unoccupied areas of the manifold? The answers to these questions could enhance the implementation of dimensional approaches for individualized neuropsychiatric care (46).

Our results must be interpreted in light of notable limitations. First, the study primarily utilized an online sample, which may not accurately represent the demographic and clinical characteristics of populations with specific mental health diagnoses. The potential differences in internet access, motivation, and the self-report nature of online studies can introduce biases that may differ from clinical settings. Consequently, the generalizability of our findings to clinical populations remains to be determined. Second, while our results are statistically robust and significant, it is important to note that the observed effect sizes are relatively small. This is not uncommon in studies of individual differences, where effect sizes often are modest due to the complex nature of human behavior and the multitude of factors influencing decision-making processes (47). Nonetheless, these small effects can still provide valuable insights into the relationships between affective states and decision-making under uncertainty. Third, our results are inherently correlational, limiting our ability to infer causal relationships between the affective states of apathy and anxiety and their impacts on decision-making processes. The observed associations provide a strong foundation for hypothesizing causal mechanisms but do not confirm them. Future studies may examine clinical samples of conditions known to affect adaptive decision-making under uncertainty, such as depression, anxiety disorders, and Parkinson’s disease, as well as interventions targeting the physiology of adaptive behavior.

## Method

### Ethics approval

The experimental procedures of all experiments were in line with the standards set by the Declaration of Helsinki and were approved by the local Research Ethics Committee of the University of Minnesota, Twin Cities. Participants provided written informed consent after the experimental procedure had been fully explained and were reminded of their right to withdraw at any time during the study.

### Participants

We recruited a sample of 1512 participants via Amazon Mechanical Turk (MTurk) and Prolific (Prolific. co); exclusion criteria included current or history of neurological and psychiatric disorders. 1001 participants completed all questionnaires and the bandit task (age range 18-54, mean ± SD = 28.446 ± 10.354 years; gender, 493 female). All participants were compensated for their time in accordance with minimum wage.

### Questionnaire measurement

Participants’ anxiety and apathy states were measured by the General Anxiety Disorder Screener (GAD-7) (36), and the Apathy-Motivation Index (35), respectively. More specifically, GAD-7 contains 7 items for assessing anxiety severity in the last two weeks. All items were rated on a 4-point scale, with higher scores indicating greater anxiety. Participants’ apathy level was measured using the 18-item Apathy-Motivation Index (AMI), which was designed to identify and measure general apathy, as well as subtypes of apathy in behavioral, social, and emotional domains. Higher scores on AMI represent greater apathy. We also measured depressive and anhedonia states by Patient Health Questionnaire (PHQ-9) (48) and Snaith-Hamilton Pleasure Scale (SHPS) (49). Our analysis did not reveal any significant results related to depressive states or anhedonia. For all questionnaire scores, see SI, Section 1 & Table S1.

### Three-armed restless bandit task

We assessed exploration-exploitation behavioral dynamics using a 300-trial three-armed restless bandit task (25). Participants were free to choose between three targets for the potential to earn a reward of 1 point. Each target is associated with a hidden reward probability that randomly and independently changes throughout the task. We seeded each participant’s reward probability walks randomly to prevent biases due to particular kinds of environments. We assessed performance by comparing the total number of rewarded trials to that expected by chance. Out of the 1001 participants, 985 accrued more rewarded trials than would be expected by chance.

### Dimensionality reduction method

Popular and valid dimensionality reduction techniques to reveal manifolds include t-distributed stochastic neighborhood embedding (t-SNE) (50), uniform manifold approximation and projection (UMAP) (37), and Principal component analysis (PCA) (51). However, t-SNE suffers from limitations, including slow computation time and loss of global data structure, and it is not a deterministic algorithm (52). The main drawback of PCA is that it is highly affected by outliers in the dataset (51). In contrast, UMAP is a deterministic and efficient algorithm, it also preserves both local and global structure of original high-dimensional data. Uniform Manifold Approximation and Projection (UMAP)

UMAP was implemented in the R language. The eight-dimensional datasets from all participants were passed into the R package *umap*, version 0.2.8.0, available at https://cran.r-project.org/web/packages/umap/) with default parameter setting as n_component = 2, n_neighbors = 15, min_dist = 0.3, metric = ‘Euclidean’. For reproducibility reasons, we fixed the random_state in this algorithm. The hyperparameter *n_neighbors* decide the radius of the search region. Larger values will include more neighbors, thus forcing this algorithm to consider more global structure of original n-dimension data. Another important hyper-parameter, *min_dist* determines the allowed minimum distance apart for cases in lower-dimensional space. *metric* defines the way that UMAP is used to measure distances along the manifold.

### Model-free analyses

We adopted some widely used model-free measures, including win-stay and lose-shift (33, 53) as the direct measurement for this learning task.

#### Win-stay

Win-stay is defined as the percentage of times that the choice in trial *t*-1 was repeated on trial *t* following a reward.

#### Lose-shift

In contrast, lose-shift equals the percentage of trials that the choice was shifted or changed when the outcome of trial *t*-1 was non-reward.

Model free results can be found at SI Section 7, Table S4.

### Mediation analyses

Mediation analysis is a statistical method used to examine the underlying mechanisms by which an independent variable influences a dependent variable through one or more mediator variables (54). In our study, we employed the bootstrapping method to estimate the mediation effect of volatility and stochasticity on the relationship between affective states (apathy and anxiety) and exploration. Bootstrapping is a nonparametric approach to effect-size estimation and hypothesis testing that is increasingly recommended for many types of analyses, including mediation (55). This method involves repeatedly resampling from the available data to generate an empirical approximation of the sampling distribution of the indirect effect (i.e., the effect of the independent variable on the dependent variable through the mediator). We used this distribution to calculate p-values and construct confidence intervals based on 5,000 resamples. Bootstrapping is preferred over other methods, such as the Sobel test because it does not assume the normality of the sampling distribution and provides more accurate confidence intervals that are bias-corrected and accelerated (54, 55). This approach offers a robust and powerful way to test mediation hypotheses, particularly in cases where the sample size is relatively small or the data violate assumptions of normality (56).

### Hidden Markov Model

We fit a Hidden Markov Model (HMM) to the behavior, to decode the hidden state of each trial for each participant. Fundamentally, the HMM has two layers, the hidden layer (i.e., state) and the observable layer. The hidden dimension should satisfy the Markov property. That is, the current hidden state only depends on the previous state but not any past model history. The observable dimension entirely depends on the current hidden states and is independent of other observations. Parameters of the hidden Markov model can be represented as *Ω* → (*T*, *O*, *c*). Specifically, *T* is the transition probabilities matrix, *O* is the observation probabilities matrix, or emissions matrix, and *c* refers to a vector with initial probabilities for each hidden state. Here, we have two hidden states, an “exploration” state and an “exploitation” state.

The transition probability between exploration and exploitation states can be represented by:

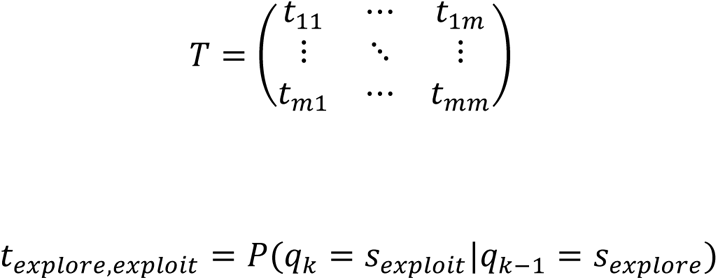

Where *t*_*explore*,*exploit*_ refers to the transition probability from hidden state *s*_*exploit*_ to another hidden state *s*_*explore*_

*k* = time instant, *m* = state sequence length

Then matrix *O* represents the transition probabilities between hidden and observable states.

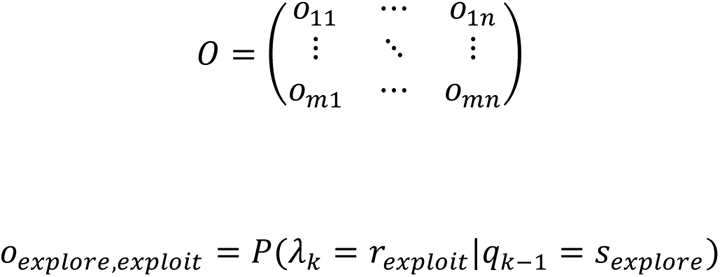

*k* = time instant, *n* = observation sequence length

Both matrix *O* and *T* satisfy the principle that the sum along the rows must be equal to 1. *c* is an m-dimensional row vector that refers to the initial probability distribution. In our current study, the initial probability was fixed and equal to the available choices.

We fit HMM via expectation-maximization using the Baum-Welch algorithm and decode hidden states from observed choice sequences by the Viterbi algorithm (57). Model results of HMM can be found at SI Section 7, Table S5.

### Kalman filter

The Kalman filter (KF) model has been widely applied in psychology and neuroscience to study various aspects of learning and decision-making (30, 31).

In the Kalman filter model for a multi-armed bandit task, *process noise* and *observation noise* refer to two distinct sources of uncertainty that affect the learning and decision-making process. Process noise represents the uncertainty in the evolution of the hidden state (reward mean) over time. It accounts for how the true state evolves from one point in time to the next. In mathematical terms, process noise is part of the state transition equation in the Kalman Filter:

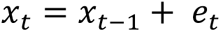

*x*_*t*_ is the state at time *t*

*e*_*t*_ is the process noise *t*, which is assumed to be drawn from a normal distribution with zeros mean and **process noise variance** *ν*. Where the *e*_*t*_∼*N*(0, *ν*).

The process noise captures the idea that the reward-means for each arm can change from one trial to the next, even in the absence of any observations. A higher process noise variance *ν* indicates a more volatile environment, where the reward means are expected to change more rapidly.

In contrast, **observation noise** represents the uncertainty in the observed rewards, given the current hidden state (reward mean). Which is assumed to be Gaussian with zero mean and a fixed variance *σ*^2^.

The observation noise captures the idea that the observed rewards are noisy and can deviate from the true reward mean due to random fluctuations or measurement errors.

A higher measurement noise variance indicates a more stochastic environment, where the observed rewards are less reliable and informative about the underlying reward means.

The Kalman Filter operates optimally when the statistical properties of the process noise and the measurement noise are accurately known.

When observation noise variance (*σ*^2^) is high relative to the process noise variance (*ν*), the Kalman gain will be small, and the model will rely more on its prior beliefs and less on noisy observations. Conversely, when the observation noise variance (*ν*), is high relative to the process noise variance (*σ*^2^), the Kalman gain will be large, and the model will update its beliefs more strongly based on the observed rewards.

### Extended Kalman filter for three-armed bandit task

The Kalman filter model can be extended to capture the effects of both volatility and stochasticity in a multi-armed bandit task (27, 58).

In the current study, process noise variance (*ν*) and observation noise variance (*σ*^2^) represent volatility and stochasticity, respectively.

A traditional assumption of the Kalman filter is that the process noise variance, *ν*, as well as the observation noise variance, *σ*^2^are constant.

Reward means update:

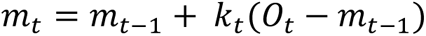

Where *m*_*t*_ is the estimated mean or value of the chosen arm at time *t*

and *O*_*t*_ is the observed reward at time *t*.

The mean update is driven by the prediction error, which is the difference between the observed reward and the previous estimate.

Kalman gain is defined as:

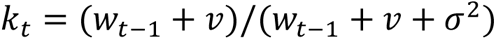

Here, *k*_*t*_ represents the Kalman gain or learning rate, which adjusts the weight given to new information based on the relative uncertainty of the prior estimate (*w*_*t*−1_) and the total noise (*ν* + *σ*^2^). When the stochasticity (*σ*^2^) is high relative to the volatility (*ν*), the Kalman gain (learning rate) will be small, and the model will rely more on its prior beliefs and less on the observations. Conversely, when the volatility (*ν*), is high relative to the stochasticity (*σ*^2^), the Kalman gain (learning rate) will be large, and the model will update its beliefs more strongly based on the observed rewards.

Variance update equation:

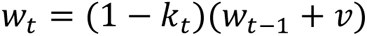

This equation updates the posterior variance (*w*_*t*_), which represents the estimate’s uncertainty after observing *O*_*t*_.

### Volatile Kalman filter for three-armed bandit task

The key difference between a standard Kalman filter and a volatile Kalman filter (VKF) is the variance of the process noise, a stochastic variable that changes with time. In other words, the VKF introduces parameters to handle the volatility in the process noise. Specifically, it allows the process noise variance *ν* to vary with the observed prediction errors, reflecting changes in environmental volatility.

Our approach here is essentially the same as that taken by Piray and Daw (27). Here, we briefly described the model details as follows.

Kalman gain:

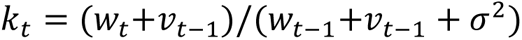

Update for the reward means:

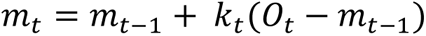

Update for posterior variance *w*_*t*_:

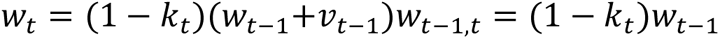

Update for volatility:

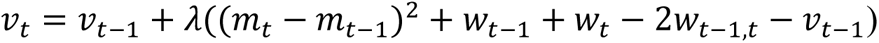

### Rescorla-Wagner models

We also fitted the data to the classical Rescorla-Wagner model. Successful adaptation in a dynamic situation requires the appropriate feedback-based learning process where individuals integrate the feedback (reward or non-reward) into the stimulus-outcome association (59). The basic reinforcement learning model, the Rescorla-Wagner model can address this process well. So the first model (RW1) was defined as:

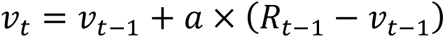

where *ν*_*t*_ is the value of the option on trial *t*.

*a* represents the general learning rate from feedback.

To verify whether participants employed distinct or shared computational responses to positive and negative feedback, we built another model with two learning rates, one for positive feedback and the other for negative feedback (33). This model (RW2) can be defined as:

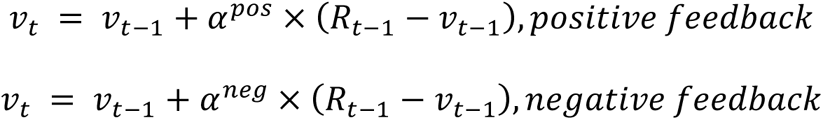

Where *v*_t_ is the value of the option on trial *t*. *α*^*pos*^and *α*^*neg*^ represent the learning rates from positive and negative feedback, respectively.

For these two models, *R*_*t*−1_ ∈ {0,1} represents the feedback received in response to participants’ choice on trial *t*-1. And *R*_*t*−1_ − *ν*_*t*−1_ represents prediction error in trial *t*-1.

We used a softmax choice function to map the value into choice. The softmax function for these four models can be defined as:

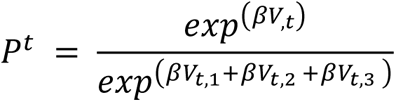

Where the *β* represents the inverse temperature with choice value.

### Model fitting and comparison

Hierarchical Bayesian inference (HBI) is a powerful method for model fitting and comparison in group studies (34). Unlike traditional approaches such as maximum likelihood estimation (MLE) or maximum a posteriori (MAP) estimation, which fit models to each subject independently, HBI simultaneously fits models to all subjects while constraining individual fits based on group-level statistics (i.e., empirical priors). This approach yields more robust and reliable parameter estimates, particularly when individual subject data is noisy or limited.

In our study, we employed HBI to fit models to choice data. The method quantifies group-level mean parameters and their corresponding hierarchical errors. To ensure that parameter estimates remain within appropriate bounds during the fitting process, we used the sigmoid function to transform parameters bounded in the unit range or with an upper bound and the exponential function to transform parameters bounded to positive values. The initial parameters of all models were obtained using a MAP procedure, with the initial prior mean and variance for all parameters set to 0 and 6.25, respectively, based on previous research (27). This initial variance allows parameters to vary widely without substantial influence from the prior.

For model comparison, we used Bayesian model selection, specifically employing the protected exceedance probability (PXP) to select the winning model. The PXP quantifies the probability that a given model is more frequent in the population than all other models under consideration while accounting for the possibility that the observed differences in model evidence may be due to chance (60). The model with the highest PXP is selected as the winning model. This approach inherently penalizes model complexity, favoring models that balance goodness-of-fit and parsimony. Model performance (log-likelihood) can be found in SI Section 8, Table 6.

### Turning point to divide the manifold into monotonically increasing and decreasing group

To divide the manifold into monotonically increasing and decreasing groups, we sorted the scores for dimension 1 in ascending order. Initially, we fitted a linear model using the first three data points located on the upper left of the manifold. We then expanded this model by sequentially including one additional data point from dimension 1, continuing this process until we incorporated the last score (i.e., the maximum dimension 1 score, situated on the upper right of the manifold). Throughout this procedure, we monitored the t-statistic of the dimension1 coefficient to assess the statistical significance of dimension1 as a predictor. Notably, a dimension 1 score of −0.671 marked the most significant negative coefficient, after which the relationship between dimension 1 and dimension2 gradually shifted to become positive (see SI Section 9, Figure S5)

### False discovery rate correction

We adopted FDR (False Discovery Rate) correction, which was introduced by Benjamini and Hochberg (61) to control the expected proportion of false positives (Type I errors).

## Supporting information

SI Section

## Acknowledgments

This work was supported by NIMH under award #R21MH127607, NIDA under award #K23DA050909, and the University of Minnesota’s MnDRIVE (Minnesota’s Discovery, Research and Innovation Economy) initiative. Natural Sciences and Engineering Council of Canada (Discovery Grant RGPIN-2020-05577), the Research Corporation for Science Advancement & Frederick Gardner Cottrell Foundation (Project #29087), and a Research Fellowship from the Jacobs Foundation to R.B.E.

We thank Iris D Vilares, A. David Redish, Cathy Chen, Seth D Koenig, and Brian Sweiss for helpful comments on the manuscript.

## Code and data availability

The source scripts used to do data analysis are published at https://github.com/hermandarrowlab/uncertainty_apathy_anxiety

Data is available upon request.

## Reference

1. A. Soltani, A. Izquierdo, Adaptive learning under expected and unexpected uncertainty. Nat. Rev. Neurosci. 20, 635–644 (2019).

2. P. Piray, N. D. Daw, A model for learning based on the joint estimation of stochasticity and volatility. Nat. Commun. 12, 6587 (2021).

3. E. Pulcu, M. Browning, The Misestimation of Uncertainty in Affective Disorders. Trends Cogn. Sci. 23, 865–875 (2019).

4. M. Husain, J. P. Roiser, Neuroscience of apathy and anhedonia: a transdiagnostic approach. Nat. Rev. Neurosci. 19, 470–484 (2018).

5. R. S. Marin, Apathy: Concept, Syndrome, Neural Mechanisms, and Treatment. Semin. Clin. Neuropsychiatry 1, 304–314 (1996).

6. F. H. Hezemans, N. Wolpe, J. B. Rowe, Apathy is associated with reduced precision of prior beliefs about action outcomes. J. Exp. Psychol. Gen. 149, 1767–1777 (2020).

7. J. Scholl, H. A. Trier, M. F. S. Rushworth, N. Kolling, The effect of apathy and compulsivity on planning and stopping in sequential decision-making. PLoS Biol. 20, e3001566 (2022).

8. Q. J. M. Huys, P. Dayan, A Bayesian formulation of behavioral control. Cognition 113, 314–328 (2009).

9. M. Browning, T. E. Behrens, G. Jocham, J. X. O’Reilly, S. J. Bishop, Anxious individuals have difficulty learning the causal statistics of aversive environments. Nat. Neurosci. 18, 590–596 (2015).

10. K. Uhr, M. J. Dugas, The role of fear of anxiety and intolerance of uncertainty in worry: an experimental manipulation. Behav Res Ther 47, 215–223 (2009).

11. C. Gagne, O. Zika, P. Dayan, S. J. Bishop, Impaired adaptation of learning to contingency volatility in internalizing psychopathology. Elife 9, e61387 (2020).

12. J. B. Hirsh, R. A. Mar, J. B. Peterson, Psychological entropy: a framework for understanding uncertainty-related anxiety. Psychol. Rev. 119, 304–320 (2012).

13. K. C. Aberg, I. Toren, R. Paz, A neural and behavioral trade-off between value and uncertainty underlies exploratory decisions in normative anxiety. Mol. Psychiatry 27, 1573–1587 (03/2022).

14. K. Witte, T. Wise, Q. J. Huys, E. Schulz, Exploring the Unexplored: Worry as a Catalyst for Exploratory Behavior in Anxiety and Depression. (2024).

15. H. Fan, S. J. Gershman, E. A. Phelps, Trait somatic anxiety is associated with reduced directed exploration and underestimation of uncertainty. Nat Hum Behav 1–12 (2022).

16. R. Smith, et al., Lower Levels of Directed Exploration and Reflective Thinking Are Associated With Greater Anxiety and Depression. Front. Psychiatry 12 (2022).

17. V. A. Holthoff, et al., Regional cerebral metabolism in early Alzheimer’s disease with clinically significant apathy or depression. Biol. Psychiatry 57, 412–421 (2005).

18. M.-C. Wen, L. L. Chan, L. C. S. Tan, E. K. Tan, Depression, anxiety, and apathy in Parkinson’s disease: insights from neuroimaging studies. Eur. J. Neurol. 23, 1001–1019 (2016).

19. D. C. Steffens, M. Fahed, K. J. Manning, L. Wang, The neurobiology of apathy in depression and neurocognitive impairment in older adults: a review of epidemiological, clinical, neuropsychological and biological research. Transl. Psychiatry 12, 1–16 (2022).

20. R. Dan, et al., Separate neural representations of depression, anxiety and apathy in Parkinson’s disease. Sci. Rep. 7, 12164 (2017).

21. S. Tinaz, et al., Distinct neural circuits are associated with subclinical neuropsychiatric symptoms in Parkinson’s disease. J. Neurol. Sci. 423, 117365 (2021).

22. C. S. Oosterwijk, C. Vriend, H. W. Berendse, Y. D. van der Werf, O. A. van den Heuvel, Anxiety in Parkinson’s disease is associated with reduced structural covariance of the striatum. J. Affect. Disord. 240, 113–120 (2018).

23. M. E. P. Seligman, Helplessness: On depression, development, and death. A series of books in psychology. 250 (1975).

24. C. S. Chen, E. Knep, A. Han, R. B. Ebitz, N. M. Grissom, Sex differences in learning from exploration. Elife 10, e69748 (2021).

25. R. B. Ebitz, E. Albarran, T. Moore, Exploration Disrupts Choice-Predictive Signals and Alters Dynamics in Prefrontal Cortex. Neuron 97, 450–461.e9 (01/2018).

26. E. A. Kaske, et al., Prolonged Physiological Stress Is Associated With a Lower Rate of Exploratory Learning That Is Compounded by Depression. Biol Psychiatry Cogn Neurosci Neuroimaging (2022). 10.1016/j.bpsc.2022.12.004.

27. P. Piray, N. D. Daw, A simple model for learning in volatile environments. PLoS Comput. Biol. 16, e1007963 (2020).

28. A. N. Hampton, P. Bossaerts, J. P. O’Doherty, The role of the ventromedial prefrontal cortex in abstract state-based inference during decision making in humans. J. Neurosci. 26, 8360–8367 (2006).

29. F. Schlagenhauf, et al., Striatal dysfunction during reversal learning in unmedicated schizophrenia patients. Neuroimage 89, 171–180 (2014).

30. S. Cheng, et al., Uncertainty-aware and multigranularity consistent constrained model for semi-supervised hashing. IEEE Trans. Circuits Syst. Video Technol. 32, 6914–6926 (2022).

31. P. Dayan, S. Kakade, P. R. Montague, Learning and selective attention. Nat. Neurosci. 3 **Suppl**, 1218–1223 (2000).

32. D. Saffron, The Rescorla - Wagner Interpretation of Blocking and Overshadowing in Pavlovian Conditioning (University of Sydney, 1979).

33. H. E. M. den Ouden, et al., Dissociable effects of dopamine and serotonin on reversal learning. Neuron 80, 1572 (2013).

34. P. Piray, A. Dezfouli, T. Heskes, M. J. Frank, N. D. Daw, Hierarchical Bayesian inference for concurrent model fitting and comparison for group studies. PLoS Comput. Biol. 15, e1007043 (2019).

35. Y.-S. Ang, P. Lockwood, M. A. J. Apps, K. Muhammed, M. Husain, Distinct Subtypes of Apathy Revealed by the Apathy Motivation Index. PLoS One 12, e0169938 (2017).

36. B. Löwe, et al., Validation and standardization of the Generalized Anxiety Disorder Screener (GAD-7) in the general population. Med. Care 46, 266–274 (2008).

37. L. McInnes, J. Healy, N. Saul, L. Großberger, UMAP: Uniform Manifold Approximation and Projection. Journal of Open Source Software 3, 861 (2018).

38. J. Aylward, et al., Altered learning under uncertainty in unmedicated mood and anxiety disorders. Nat Hum Behav 3, 1116–1123 (2019).

39. D. W. Grupe, J. B. Nitschke, Uncertainty and anticipation in anxiety: an integrated neurobiological and psychological perspective. Nat. Rev. Neurosci. 14, 488–501 (2013).

40. M. Fahed, D. C. Steffens, Apathy: Neurobiology, assessment and treatment. Clin. Psychopharmacol. Neurosci. 19, 181–189 (2021).

41. J. Pagonabarraga, J. Kulisevsky, A. P. Strafella, P. Krack, Apathy in Parkinson’s disease: clinical features, neural substrates, diagnosis, and treatment. Lancet Neurol. 14, 518–531 (2015).

42. M. Pessiglione, F. Vinckier, S. Bouret, J. Daunizeau, R. Le Bouc, Why not try harder? Computational approach to motivation deficits in neuro-psychiatric diseases. Brain 141, 629–650 (2018).

43. J. S. Beck, Cognitive behavior therapy: Basics and beyond (Guilford Press, 2011).

44. M. G. Craske, J. L. Mystkowski, “Exposure therapy and extinction: Clinical studies” in Fear and Learning: From Basic Processes to Clinical Implications, (American Psychological Association, 2007), pp. 217–233.

45. G. S. Alexopoulos, P. Arean, A model for streamlining psychotherapy in the RDoC era: the example of “Engage.” Mol. Psychiatry 19, 14–19 (2014).

46. B. N. Cuthbert, T. R. Insel, Toward the future of psychiatric diagnosis: the seven pillars of RDoC. BMC Med. 11, 126 (2013).

47. D. C. Funder, D. J. Ozer, Evaluating effect size in psychological research: Sense and nonsense. Adv. Methods Pract. Psychol. Sci. 2, 156–168 (2019).

48. K. Kroenke, R. L. Spitzer, J. B. W. Williams, The PHQ-9. J. Gen. Intern. Med. 16, 606–613 (2001).

49. R. P. Snaith, et al., A scale for the assessment of hedonic tone the Snaith-Hamilton Pleasure Scale. Br. J. Psychiatry 167, 99–103 (1995).

50. G. E. Hinton, S. Roweis, Stochastic neighbor embedding. Adv. Neural Inf. Process. Syst. 15 (2002).

51. I. T. Jolliffe, J. Cadima, Principal component analysis: a review and recent developments. Philos. Trans. A Math. Phys. Eng. Sci. 374, 20150202 (2016).

52. H. I. Rhys, Machine Learning with R, the tidyverse, and mlr (Simon and Schuster, 2020).

53. A. Bari, et al., Serotonin modulates sensitivity to reward and negative feedback in a probabilistic reversal learning task in rats. Neuropsychopharmacology 35, 1290–1301 (2010).

54. A. F. Hayes, Introduction to mediation, moderation, and conditional process analysis: a regression-based approach. xvii (Guilford Press, 2017).

55. D. P. Mackinnon, C. M. Lockwood, J. Williams, Confidence Limits for the Indirect Effect: Distribution of the Product and Resampling Methods. Multivariate Behav. Res. 39, 99 (2004).

56. K. J. Preacher, A. F. Hayes, Asymptotic and resampling strategies for assessing and comparing indirect effects in multiple mediator models. Behav. Res. Methods 40, 879–891 (2008).

57. J. P. Coelho, T. M. Pinho, J. Boaventura-Cunha, Hidden Markov Models: Theory and Implementation using Matlab® (CRC Press, 2019).

58. K. Chakroun, D. Mathar, A. Wiehler, F. Ganzer, J. Peters, Dopaminergic modulation of the exploration/exploitation trade-off in human decision-making. Elife 9 (2020).

59. B. U. Forstmann, R. Ratcliff, E.-J. Wagenmakers, Sequential sampling models in cognitive neuroscience: Advantages, applications, and extensions. Annu. Rev. Psychol. 67, 641–666 (2016).

60. L. Rigoux, K. E. Stephan, K. J. Friston, J. Daunizeau, Bayesian model selection for group studies — Revisited. Neuroimage 84, 971–985 (2014).

61. Y. Benjamini, Y. Hochberg, Controlling the false discovery rate: A practical and powerful approach to multiple testing. J. R. Stat. Soc. 57, 289–300 (1995).

