## Supplementary material for "Distinct computational mechanisms of uncertainty processing explain opposing exploratory behaviors in anxiety and apathy": SI Section

### Individual differences in uncertainty evaluation explain opposing exploratory behaviors in anxiety and apathy

**This PDF file includes:**

Supporting text

9 SI Sections

Figure. S1 to S5

Tables S1 to S6

**Section1. Descriptive statistic for all questionnaires in current study**

We reported the basic statistical information for all questionnaires in the current study in Table S1.

Table S1. Descriptive statistics for all questionnaires.

| | PHQ-9 | GAD-7 | SHPS | Apathy | Apathy-ES | Apathy-BA | Apathy-SM |
|---|---|---|---|---|---|---|---|
| Mean | 9.53 | 7.17 | 1.92 | 30.46 | 10.54 | 12.69 | 7.23 |
| SD | 6.13 | 5.54 | 2.26 | 9.32 | 5.04 | 4.80 | 4.22 |
| Minimum | 0 | 0 | 0 | 4 | 0 | 0 | 0 |
| Maximum | 28 | 21 | 14 | 64 | 24 | 24 | 24 |

[a]PHQ-9 = Patient Health Questionnaire; SHPS = Snaith-Hamilton Pleasure Scale; GAD-7 = General Anxiety Disorder Screener; Apathy-BA = Apathy behavioral activation; Apathy-ES = Apathy emotional sensitivity; Apathy-SM = Apathy social motivation.

**Section 2. Distinct computational processes between high anxious and high apathetic individuals.**

Furthermore, comparisons showed that high anxious individuals had higher volatility estimates than those with high apathy (t(449) = 2.75, p=0.006). In contrast, high apathetic individuals had higher stochasticity estimates than their high anxiety counterparts (t(449) = -3.01, p=0.002), resulting in a higher learning rate among the high anxiety group (t(449) = 3.04, p=0.002) (Figure S1).

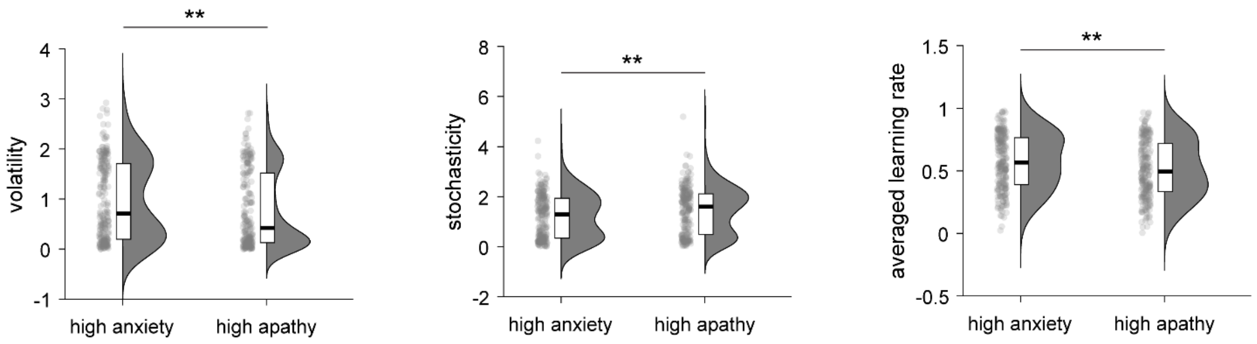


**Figure S1.** Distinct computational processes between highly anxious and highly apathetic individuals. The high anxiety group showed higher volatility, lower stochasticity, and a higher learning rate than the high apathy group. ***p*<0.01;

**Section 3. Mediation model**

We conducted a mediation analysis using anxiety, the exploration after non-reward feedback (*P(explore | 0)*), and the $v/s$. The results demonstrate that the relationship between anxiety and the tendency to switch after receiving no reward is significantly mediated by the $v/s$ (Figure S2).

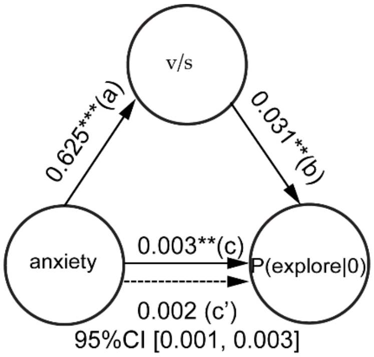


**Figure S2.** The ratio of volatility to stochasticity significantly partially mediated the relationship between anxiety and exploration after undesired feedback. $^{**}p<0.01$; $^{***}p<0.001$;

**Section 4. The robust low-dimensional space**

We conducted t-Distributed Stochastic Neighbor Embedding (t-SNE) and Principal Component Analysis (PCA) to confirm the manifold. The same eight-dimensional datasets from all participants were passed into the R package Rtsne, version 0.16 (available at https://cran.r-project.org/web/packages/Rtsne/index.html) with default parameter setting as n_component = 2, perplexity = 30, min_iter = 1000, metric = 'Euclidean'. We showed a similar manifold shape as UMAP found. Like in the main text, we also mapped model-free indices, as well as parameters from HMM onto t-SNE manifolds. The meaning of gradient change here is the same as with the UMAP manifold.

The low-dimensional space from PCA is also quite similar to the manifold from UMAP and t-SNE. All other correlation results based on t-SNE and PCA scores can be found in the tables below.

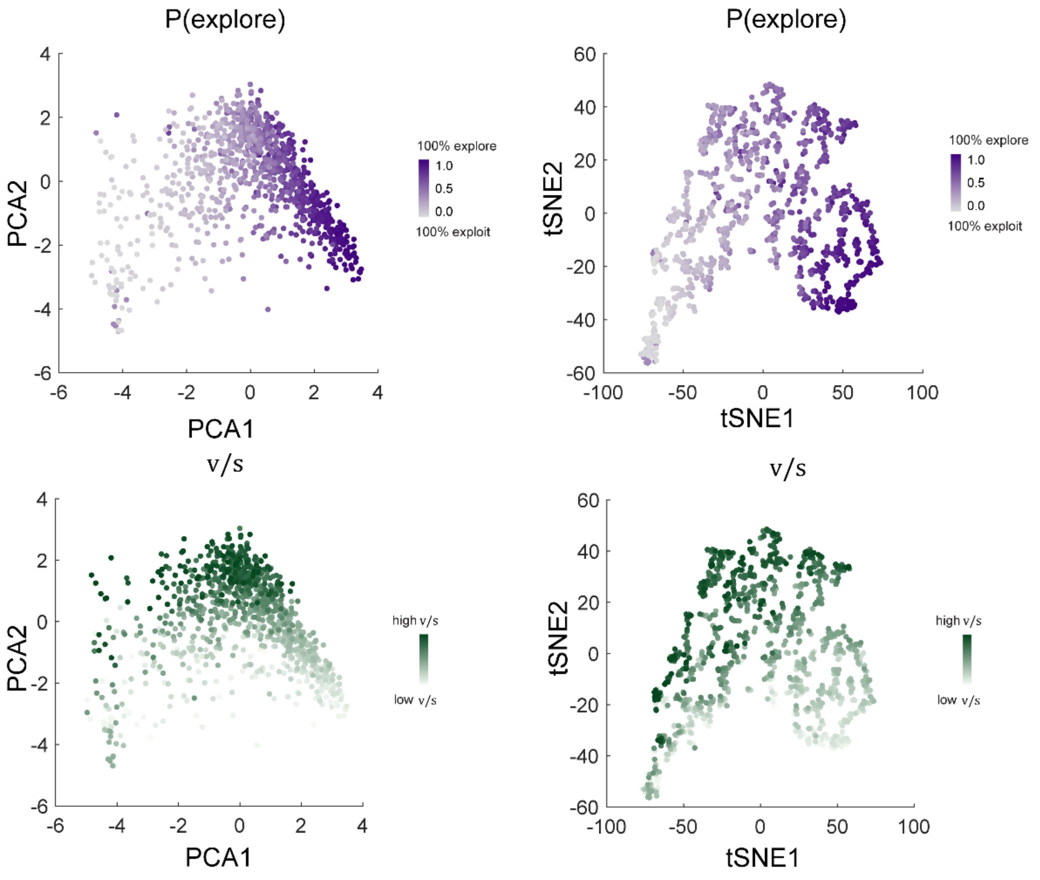


**Figure S3.** The low-dimensional space from PCA and tSNE, and the representation for P(explore), the ratio of volatility to stochasticity.

**Table S2**

| | t-SNE1 | t-SNE2 | PCA1 | PCA2 |
|---|---|---|---|---|
| Switch% | -0.914*** | -0.323*** | -0.967*** | -0.139*** |
| Stay% | 0.914*** | 0.323*** | 0.967*** | 0.139*** |
| Win.stay | 0.713*** | 0.608*** | 0.832*** | 0.513*** |
| Lose.shift | -0.631*** | 0.482*** | -0.665*** | 0.603*** |
| Exploration % | -0.912*** | -0.250*** | -0.937*** | -0.016*** |
| Exploitation % | 0.912*** | 0.250*** | 0.937*** | 0.016 |
| Exploit-exploit | 0.536*** | 0.434*** | 0.683*** | 0.357*** |
| Explore-explore | -0.484*** | 0.111*** | -0.449*** | 0.199*** |
| Explore-exploit | 0.484*** | -0.111*** | 0.449*** | -0.199*** |
| Exploit-explore | -0.536*** | -0.434*** | -0.683*** | -0.357*** |

| | | | | |
|---|---|---|---|---|
| **v/s** | -0.179*** | 0.716** | -0.168*** | 0.832*** |

[a]all significant P-values reported here survive FDR correction.

(Original Benjamini & Hochberg FDR procedure, p<0.05)

**Section 5. No quadratic relationship between apathy, anxiety, and exploration**

There is no significant quadratic relationship between apathy, anxiety, and exploration, neither between these affective states nor the ratio of volatility to stochasticity.

**Table S3.**

| **Linear models** | $\textbf{stats}_{\textbf{quadratic, apathy}}$ | $\textbf{stats}_{\textbf{quadratic, anxiety}}$ |
|---|---|---|
| P(switch)~1+ apathy+anxiety+anxiety$^2$+apathy$^2$ | $p=0.430$ | $p=0.191$ |
| P(exploration)~1+ apathy+anxiety+anxiety$^2$+apathy$^2$ | $p=0.354$ | $p=0.087$ |
| v/s ~1+ apathy+anxiety+anxiety$^2$+apathy$^2$ | $p=0.209$ | $p=0.746$ |

**Section 6. Non-linear relationship between the ratio of volatility to stochasticity and exploration.**

The results revealed that both the linear and quadratic terms are significant (linear term, coefficient = 0.02, SE = 0.003, $t(996)=5.83$, $p<10^{-9}$; quadratic term, coefficient = 0.005, SE = $7.86\times10^{-4}$, $t(996)=6.59$, $p<10^{-11}$), indicating a complex, non-linear relationship between the ratio of volatility to stochasticity and exploration.

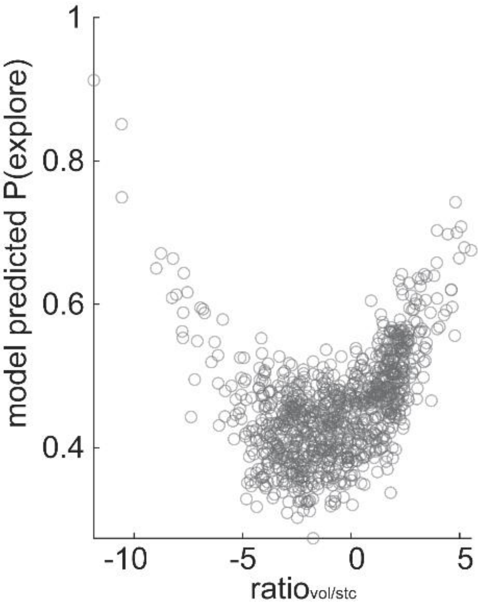


**Figure S4.** Linear and quadratic relationship between Ratio of volatility to stochasticity and exploration.

**Section 7. Model-free and model-based results**

**Table S4**

| | Model-free indices | | | |
|---|---|---|---|---|
| | Stay % | Switch % | Win. Stay | Lose.shift |
| Mean | 0.646 | 0.353 | 0.861 | 0.732 |
| SD | 0.180 | 0.180 | 0.210 | 0.204 |

**Table S5**

| | Model-based indices from HMM | | | | | |
|---|---|---|---|---|---|---|
| | Exploitation% | Exploration % | Exploit-Exploit | Exploit-Explore | Explore-Explore | Explore-Exploit |
| Mean | 0.544 | 0.455 | 0.811 | 0.188 | 0.818 | 0.181 |
| SD | 0.243 | 0.243 | 0.215 | 0.215 | 0.146 | 0.146 |

**Section 8. Model performance**

**Table 6**

| All models | loglikelihood |
|---|---|
| RW1 | -2.1456e+05 |
| RW2 | -2.1378e+05 |
| **KF** | **-1.7346e+05** |
| VKF | -1.8786e+05 |

**Section 9. Turning point to divide the manifold into monotonically increasing and decreasing group**

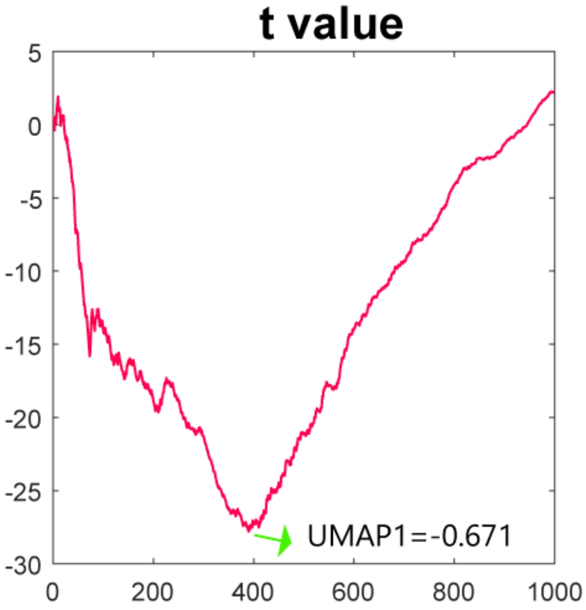


**Figure S5.** A dimension 1 score of -0.671 marked the most significant negative coefficient, after which the relationship between dimension 1 and dimension2 gradually shifted to become positive
